# Cases of Cutaneous Leishmaniasis in a peri-urban settlement in Kenya, 2016

**DOI:** 10.1101/536557

**Authors:** Isaac Ngere, Waqo Gofu, Abdikadir Isack, Joshua Muiruri, Mark Obonyo, Sultani Matendechero, Zeinab Gura

**Affiliations:** Washington State University Global Health Program Kenya; Kenya Field Epidemiology and Laboratory Training Program; Ministry of Health, Republic of Kenya

**Keywords:** Cutaneous Leishmaniasis, Kenya, forests

## Abstract

**Background:** Cutaneous Leishmaniasis is a neglected tropical disease caused by a protozoan and transmitted by sand-fly bite. Following reports of a possible outbreak of cutaneous leishmaniasis in 2016, we conducted a review of hospital records and a follow up case control study to determine the magnitude of the disease, characterize the cases and identify factors associated with the disease in Gilgil, a peri-urban settlement in Central Kenya.

**Methods:** We reviewed hospital records, conducted active case search in the community and carried out a case-control study. Medical officers in the study team made clinical diagnosis of cutaneous leishmaniasis cases based on presence of a typical skin ulcer. We enrolled 58 cases matched by age and residence to 116 controls in a case control study. We administered structured questionnaires and recorded environmental observations around homes of cases and controls. Simple proportions, means and medians were calculated for categorical data and continuous data respectively. Logistic regression models were constructed for individual, indoor and outdoor factors associated with the outbreak.

**Results:** We identified 255 suspected cases and one death; Females constituted 56% (142/255), median age of the cases was 7 years (IQR 14). Cases were clustered around Gitare (28.6%, 73/255) and Kambi-Turkana (14%, 36/255) with seasonal peaks between June-November. Among individual factors, staying outside the residence in the evening after sunset (OR 4.1, CI 1.2-16.2) and occupation involving visiting forests (OR 4.56, CI 2.04-10.22) had significant associations with disease. Sharing residence with a cutaneous leishmaniasis patient (OR 14.4, CI 3.8-79.3), a house with alternative roofing materials (OR 7.9, CI 1.9-45.7) and residing in a house with cracked walls (OR 2.3, CI 1.0-4.9) were significant among indoor factors while sighting rock hyraxes near residence (OR 5.3, CI 2.2-12.7), residing near a forest (OR 7.8, CI 2.8-26.4) and living close to a neighbour with cutaneous leishmaniasis (OR 6.8, CI 2.8-16.0) had increased likelihood of disease. Having a cultivated crop farm surrounding the residence (OR 0.1, CI 0.0-0.4) was protective.

**Conclusions/Significance:** This study reveals the large burden of cutaneous leishmaniasis in Gilgil. There is strong evidence for both indoor and outdoor patterns of disease transmission. Occupations and activities that involve visiting forests or residing near forests and sharing a house or neighbourhood with a person with CL were identified as significant exposures of the disease. The role of environmental factors and wild mammals in disease transmission should be investigated further

**Author summary:** Leishmaniasis is a group of diseases caused by a protozoa (*Leishmania)* and affects humans and other mammals following the bite of an infected sand-fly. Cutaneous form of the disease (cutaneous leishmaniasis) is considered a neglected tropical disease mainly affecting the poor destabilized or migrant populations in rural areas. Recently, the disease has expanded its geographical range and invaded previously non-endemic areas including areas surrounding large urban centres that are experiencing human population influx leading to multiple localised disease outbreaks. In this paper, we report findings of a study we conducted to determine the burden and factors promoting the spread of cutaneous leishmaniasis in a peri-urban settlement in Kenya. Our results indicate a high burden of cutaneous leishmaniasis in this area and an association of the disease with several groups of factors at individual, indoor and outdoor environments. Many cases of cutaneous leishmaniasis were linked to activities that involved visiting the forested areas around homes, underpinning the significance of human activity in forests in these areas in spread of the disease.

## Introduction

Leishmaniasis is a disease caused by a protozoan, *Leishmania* and is transmitted to humans and other mammals by the bite of a female phlebotomite sand-fly. Three forms of the disease affect humans; cutaneous, muco-cutaneous and visceral forms (*Kala-azar*). The disease is considered a neglected tropical disease mainly affecting the rural poor. Cutaneous leishmaniasis commonly occurs in clusters among destabilized or migrant populations in low socio-economic set ups with the current trend in distribution of new infections indicating a progressive spread of the disease to previously non-endemic areas (1–5).

Worldwide, over 350 million people are estimated to be at risk of cutaneous leishmaniasis and 2 million new infections are reported annually (6). Owing to challenges in surveillance and reporting, the burden of cutaneous leishmaniasis is grossly underestimated (4). Though disfiguring and debilitating in the affected people, the disease is rarely fatal, hence little attention has been given to prevention and control measures by health authorities (4,5). Proven control strategies include vector eradication and early treatment of insect bites in endemic areas. Insect vector control activities such as indoor insecticide residual sprays, insecticide impregnated barriers (bed nets, curtains, clothes, carpets), environmental spraying, and control of intermediate hosts (rodents) are effective but expensive when rolled out on a large scale (4). Therefore, targeted control programs guided by an understanding of local drivers of the disease including lifestyle and environmental factors would not only ensure success of such programs but will also provide significant cost savings and value for money (5,7).

Recurrent outbreaks of cutaneous leishmaniasis in Kenya, Ethiopia and South Sudan have been reported in the past and are often associated with high morbidity. Kenya is classified by WHO as endemic for cutaneous leishmaniasis. However, there is relative scarcity of published data on the extent, burden and risk factors of cutaneous leishmaniasis (8). In other parts of the world, urbanization and expansion of farming and other human activities into forests is often associated with disease outbreaks (5). In Kenya, areas around the Rift Valley escarpments and major mountains are known natural habitats for sand-flies (9–11). So far, more than 48 species of sand-flies, including the vector for CL, have been identified in various habitats in Kenya (12). Recently, the areas around the Rift valley in Kenya have been experiencing increased population and environmental pressure from in-migration and increased human activities in forests (13,14).

In early 2016, the health ministry of Kenya received notification about increase in cases of a skin disease suspected to be cutaneous leishmaniasis in Nakuru County, in south-eastern rift valley. We report the findings of a records review and a follow up case control study we conducted to determine the magnitude of the disease, characterize the cases and identify factors associated with the disease.

## Methods

### Study site

We conducted the study in Gilgil Sub-county, a rapidly growing peri-urban settlement located in south-eastern part of the Great Rift Valley in Kenya, between 20^th^ January and 3^rd^ February 2016 **(Fig. 1).** The terrain of Gilgil sub county is generally mountainous to the north with plenty of rocky escarpments. The southern end of the sub county is mainly composed of undulating plains, flat grazing lands and solidified larva forming large crevices and rocky caves with wild mammals and rodents (9,10). Scattered housing characterizes this new settlement and lately the area has experienced a surge in population since the region is regarded as high potential yet cheaper alternative to the town life in Nairobi city or Nakuru town (14). The sub county is traversed by the busy Nairobi-Eldoret highway and is a preferred destination for potential peri-urban home-owners due to its ease of access. Previous studies in this area have identified the insect vector (*Sand-fly)* and the agent (*Leishmania Donovani, Leishmania infantum* and *Leishmania Chagas*) (9–11)

**Figure 1:**
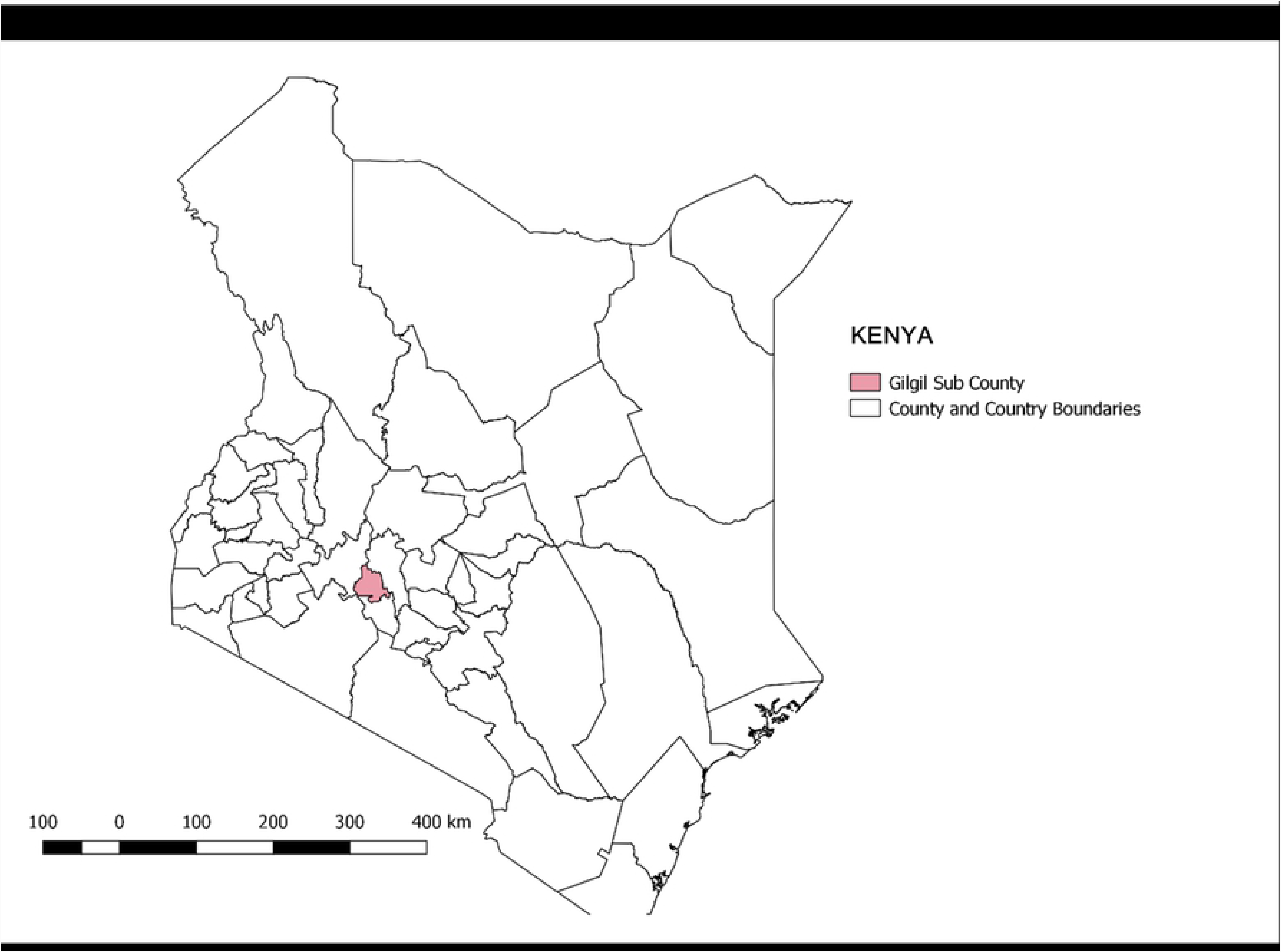
Map of the Investigation area, Gilgil, Kenya 2016. This map was drawn on QGIS Version 2.18.15 using mapping resources from International Livestock Research Institute (ILRI) (15,16)

### Study design

We reviewed hospital records and conducted door to door search to identify cases. Cases were then enrolled in a population-based case control study.

### Review of records

We used a standard data abstraction tool to review records covering nine health facilities in Morendat and Mbaruk wards. We included entries made in outpatient, inpatient, laboratory, and specialist clinic registers between January 2010 and January 2016.

A suspected case of cutaneous leishmaniasis was defined based on clinical diagnosis recorded in hospital records as ‘skin ulcer’, ‘skin wound’, ‘plaque’, ‘dermatitis’, ‘skin infection’ or ‘cutaneous leishmaniasis’. A probable case was defined as a resident with a typical ulcer or plaque (typical raised edges and depressed centre) ascertained by a medical officer in the study team during the study period. Due to logistical challenges, no laboratory confirmation for cutaneous leishmaniasis was done on the suspected or the probable cases. All entries that matched suspected, probable or confirmed case definitions were included in the study line list.

We also abstracted information such as name, sex, age, date seen at the facility, residence, signs and symptoms, diagnosis and treatment given.

### Enrolment of cases and controls into a Case Control study

To determine the risk factors of cutaneous leishmaniasis infection in the study population, we conducted a follow-up case-control study. Cases and controls consisted of eligible residents found in the study area during the study period (20^th^ January-3^rd^ February).

#### Sample size

Using OpenEpi, we calculated a sample size of 174 (2 controls per case) to be able to achieve a power of at least 80% at the 5% significance level, able to detect an odds ratio (OR) of ≤0.3 for an exposure present in 31% of controls (17–19). The exposure chosen was use of mosquito nets.

#### Case and control recruitment

We recruited 59 cases for the case control study: 41 cases were selected from the line list developed in the records review and a further 18 cases were identified during active house-to-house survey **(Fig. 2).** The investigation team comprising a field epidemiologist, 2 medical doctors, a laboratory scientist and 2 public health specialists, worked with community-based locators who included community health volunteers and local chiefs to locate and identify cases for inclusion in the study. In each village, the number of cases that were recruited in the case control study was allocated by probability proportional to size sampling based on the proportion of residents from that village with suspected cutaneous leishmaniasis from the line list.

**Figure 2:**
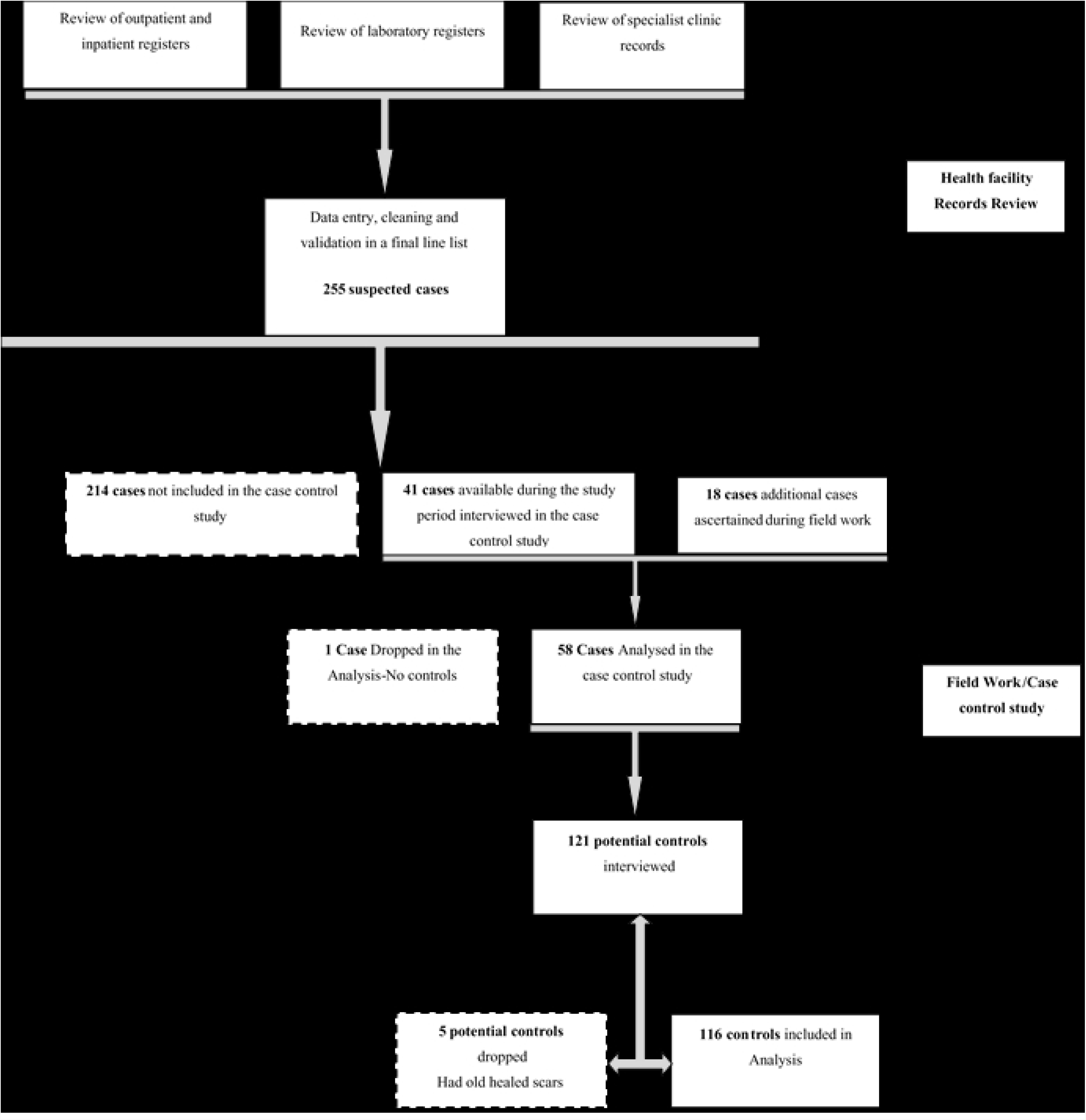
Flow diagram of selection of cases and controls before and after field work. We attempted to match each case to two community-based controls by age using the following criteria: Cases less than two years of age were matched to controls within two years, cases 2–4 years old were matched to controls within 3 years, cases 5–19 years to controls within 5 years, cases 20–59 year’s old to controls within10 years, and cases more than 60 years old to controls within 20 years. One case was dropped in the final analysis owing to lack of suitable controls **(Fig. 2).**

Controls were selected from among residents of the same age group and living in the same or neighbouring village(s) as the case patients, and had no typical ulcer, wound or scar upon inquiry and examination by medical doctors in the study team. To locate a possible control, a member of the investigation team would spin a bottle while standing afront the case’s household to determine the direction of movement. A random number between 2 and 5 was drawn to indicate the number of houses in the chosen direction to be passed before the team would attempt to recruit a control. This process was repeated until two eligible controls were recruited for each case.

#### Data collection and analysis

A structured questionnaire was administered to cases and controls through face to face interviews. We collected information on demographic profile, clinical presentation, risk factor profile and environmental exposures among cases and controls regarding cutaneous leishmaniasis. Exposure data in both cases and controls was collected in relation to the year of onset of symptoms in the case patients. Environmental observations around the home were made by the study personnel and recorded in the questionnaires.

All data from the questionnaires were entered and cleaned using Epi info version 7(CDC, Atlanta GA, USA) and Microsoft Excel ™ (Microsoft, Seattle, WA, USA). Simple proportions, means and medians were calculated for categorical data and continuous data respectively. To identify factors associated with the outbreak, chi square tests were computed for categorical data, and odds ratios and 95% CI calculated. Risk factors were categorised into three groups in the analysis: individual, indoor and outdoor factors. Factors with a *P≤0.05* in the bivariate analysis were considered significant and were included in the ‘group model’ for each category of factors. To develop the final model, regression analysis was conducted using backward elimination method, starting with all factors that had a P≤0.2 at the in the group model to determine independent risk factors.

### Human subject’s protection

This study was conducted as part of Ministry of Health (MOH) led public health response to an acute event and as such did not require a review by an ethical review body. Oral consent was obtained from the case-control study subjects and documented in the study questionnaires. A study protocol was developed and authorisation to conduct the study was given by the Kenyan Ministry of Health through the Field Epidemiology and Laboratory Training program (K-FELTP), Nakuru county and respective health facilities where recorded were abstracted. Permission was also sought from the county department of health, Nakuru County. Measures were taken to assure confidentiality of the information provided during these interviews. Review of surveillance data was conducted as part of routine surveillance by the MOH, and no personal identifying information was collected or analysed. Measures were taken to ensure collected data were properly stored and secured in both paper and electronic databases. Individuals who were found to have active lesions at the time of the study were referred for free treatment at Nakuru County Referral Hospital

## Results

### Review of records

From the review of health facility records and house-to-house survey, we identified a total of 255 cases and one death suspected to be due to cutaneous leishmaniasis between 2010-2016. Of the identified cases, females constituted 48.6% (124/255) and the median age was 7 years (IQR 14). Cases were spread throughout the study area, with most cases (28.6% or 73/255) originating from *Gitare* village and 14.1% (36/255) from *Kambi Turkana* village **(Table 1).**

**Table 1:**
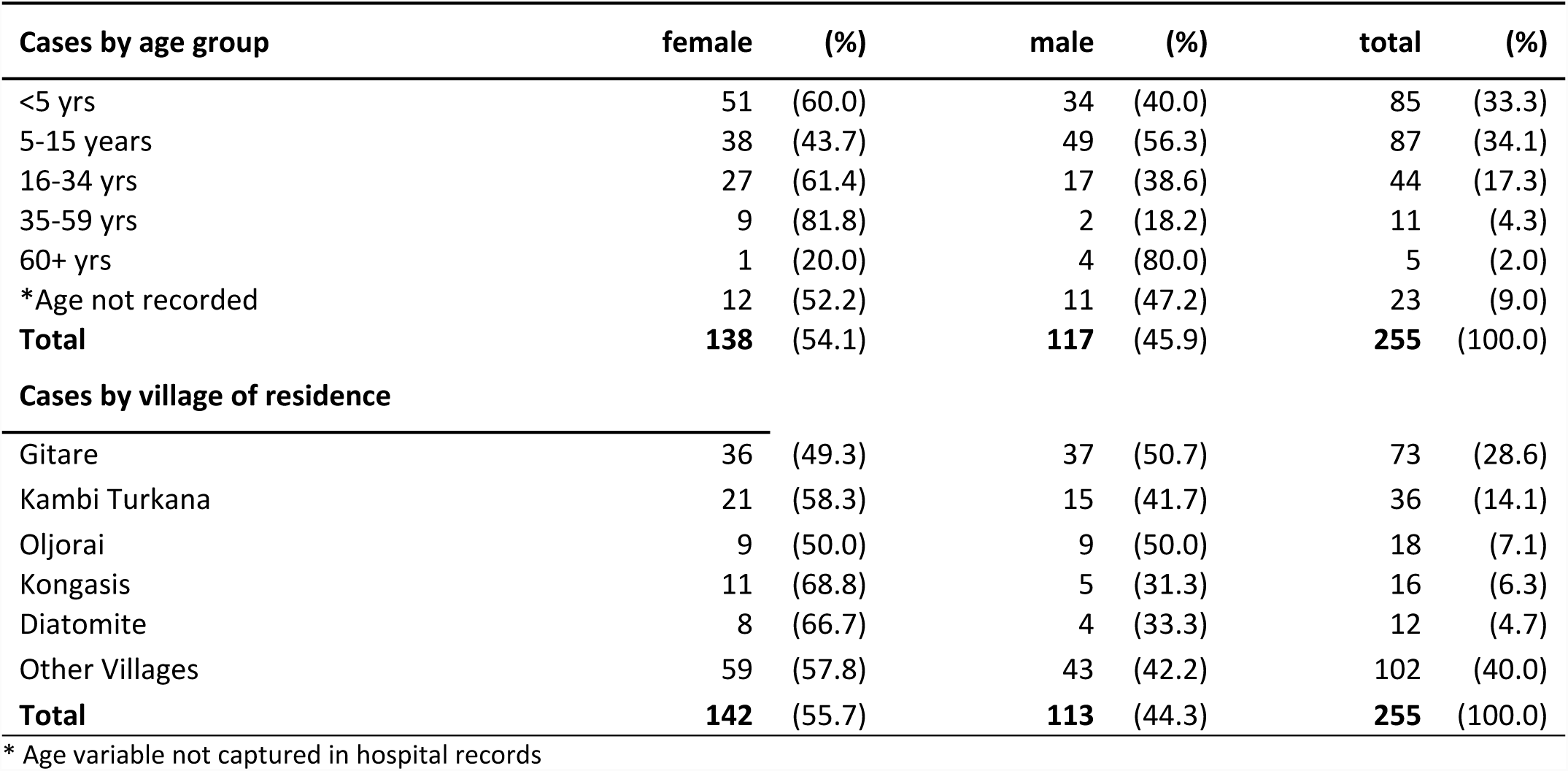
Demographic characteristics of suspected cases of Cutaneous Leishmaniasis based on records review in Gilgil, 2010-2016 (n=255)

| Cases by age group | female | (%) | male | (%) | total | (%) |
| --- | --- | --- | --- | --- | --- | --- |
| <5 yrs | 51 | (60.0) | 34 | (40.0) | 85 | (33.3) |
| 5-15 years | 38 | (43.7) | 49 | (56.3) | 87 | (34.1) |
| 16-34 yrs | 27 | (61.4) | 17 | (38.6) | 44 | (17.3) |
| 35-59 yrs | 9 | (81.8) | 2 | (18.2) | 11 | (4.3) |
| 60+ yrs | 1 | (20.0) | 4 | (80.0) | 5 | (2.0) |
| *Age not recorded | 12 | (52.2) | 11 | (47.2) | 23 | (9.0) |
| **Total** | **138** | **(54.1)** | **117** | **(45.9)** | **255** | **(100.0)** |
| **Cases by village of residence** |  |  |  |  |  |  |
| Gitare | 36 | (49.3) | 37 | (50.7) | 73 | (28.6) |
| Kambi Turkana | 21 | (58.3) | 15 | (41.7) | 36 | (14.1) |
| Oljorai | 9 | (50.0) | 9 | (50.0) | 18 | (7.1) |
| Kongasis | 11 | (68.8) | 5 | (31.3) | 16 | (6.3) |
| Diatomite | 8 | (66.7) | 4 | (33.3) | 12 | (4.7) |
| Other Villages | 59 | (57.8) | 43 | (42.2) | 102 | (40.0) |
| **Total** | **142** | **(55.7)** | **113** | **(44.3)** | **255** | **(100.0)** |

\* Age variable not captured in hospital records

Cases of suspected cutaneous leishmaniasis were recorded throughout the period between 2010 and 2016 with the majority of cases being recorded in the second half of each year (June-November). Most (23.5%) cases of suspected leishmaniasis were recorded in 2014 while the least (11.8%) cases were recorded in 2011. **(Fig. 3)**

**Figure 3:**
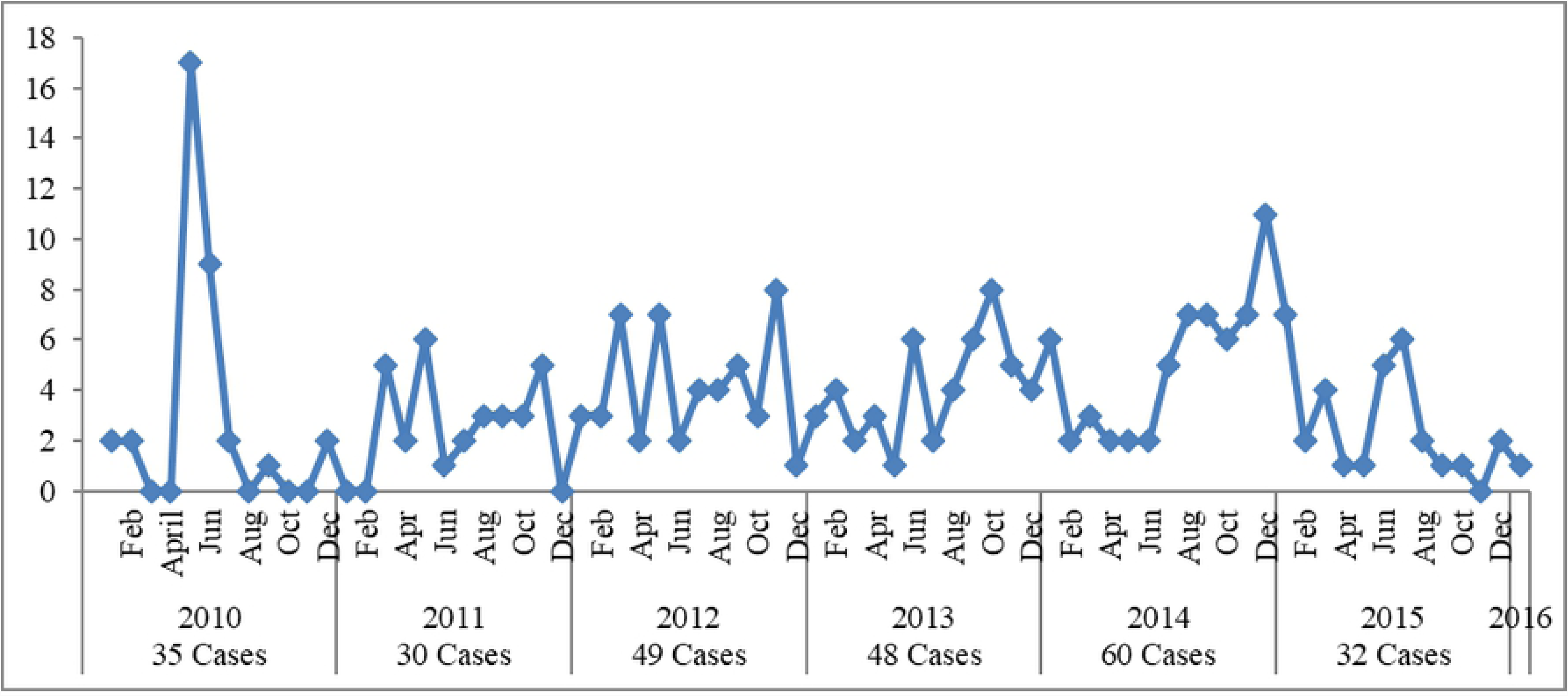
Trends of patients treated for suspected cutaneous leishmaniasis in health facilities in Gilgil between January 2010 and January 2016 (N=255)

### Case control study

**Table 2** summarizes the demographic characteristics of cases and controls enrolled in the study. Males constituted 55.2% (32/58) of the cases and 49.1% (57/116) of the controls. Cases were predominantly young persons aged below 18 years (58.6%) or older persons aged 35 years and above (32.8%). The youngest case was 2 years while the oldest was 86 years old. As expected, there were no differences in age distribution between cases and controls. The distribution of other social and demographic variables studied among cases and controls was comparable between cases and controls.

**Table 2:**
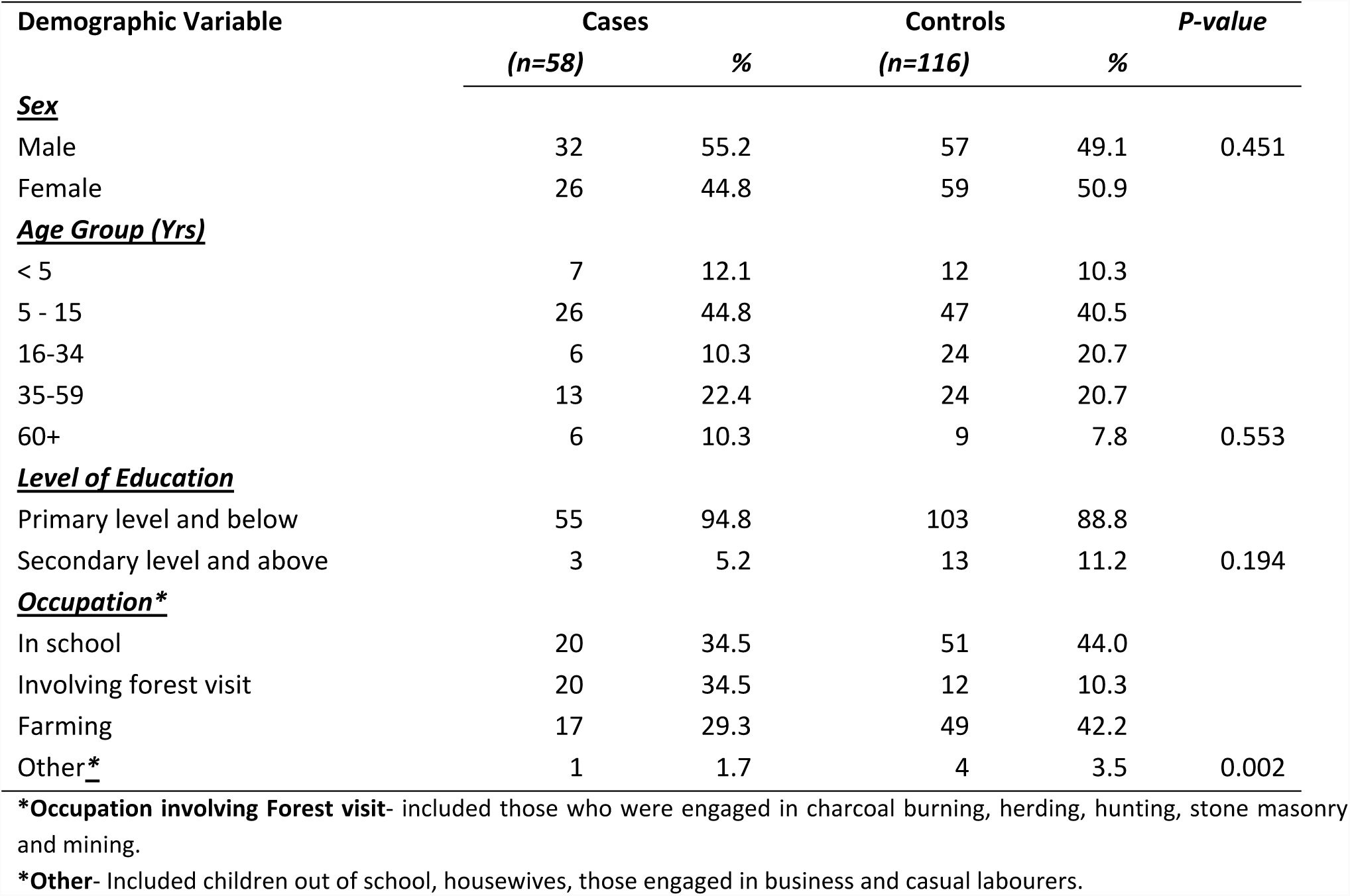
Demographic characteristics of cases and controls in Gilgil, Kenya 2016.

Among the cases, the median duration of illness was 2 years (range 1-4 years). All the cases had typical ulcers, while 84.5% (49/58) of the cases had a papule/nodule. A majority of lesions were located on the head and neck regions (81.6%, or 40/49) while 6.1% (3/49) were in the hands and 2.0% (1/49) were located in the foot. In thirteen cases (22.4%), both active ulcerative lesions and scars were found. Other symptoms observed among cases included pruiritus (15.5%), rash (5.2%), bruising (5.2%), skin infections (5.2%) and nasal stuffiness (3.4%). Various wound treatment remedies were cited by the cases including herbal medication (72.1%), over-the-counter skin ointments (41.9%) and antibiotics (25.6%).

In terms of household ownership and use of mosquito-nets, 10.3% (6/58) of the cases and 18.1% (21/116) of the controls owned and slept under a mosquito-net every night. Household indoor residual spraying (IRS) was reported by less than 5% of both cases and controls. Sighting of wild animals around homes was common among both cases and controls: Some 87.9% of cases and 57.8% of controls reported sighting rock hyraxes and 75.9% of the cases and 69.0% of the controls reported sighting mongooses near their homes. In terms of housing condition, 82.8% of cases and 97.4% of the controls lived in houses with corrugated iron sheet roofing, 56.9% of cases and 59.5% of the controls lived in houses with earthen floor, 27.6% of cases and 21.6% of the controls lived in houses with cracked floors. The majority of cases (91.4%) lived in houses located near a forest compared to 57.8% controls.

### Risk factor analysis

Potential risk factors were categorised into three groups for purposes of analysis: factors related to the individual, factors related to indoor dwelling environment and factors related to outdoor environment. Table 3 shows the distribution of cases and controls with corresponding crude and adjusted odds ratios and for the variables analysed in the 3 ‘group models’.

**Table 3:** Analysis of the risk factors associated with suspected cutaneous leishmaniasis in Gilgil, 2016.

| Variable | Cases<br>N=58 |  | Controls<br>N=116 |  | Crude OR (95% CI) | Adjusted OR<br>(95% CI) | P value |
| --- | --- | --- | --- | --- | --- | --- | --- |
|  | n | % | n | % |  |  |  |
| **A. Factors related to the individual** |  |  |  |  |  |  |  |
| Male Sex | 32 | 55.2 | 57 | 49.1 | 1.3 (0.7-2.4) | 1.0 (0.5-1.9) | 0.892 |
| Education Level: Primary and below | 55 | 94.8 | 103 | 88.8 | 2.3 (0.6- 13.1) | 2.5 (0.6-9.6) | 0.190 |
| Spending time outside home after sunset | 9 | 15.5 | 5 | 4.3 | 4.1 (1.2- 16.2) | 2.1 (0.6-7.7) | 0.284 |
| Net use by the individual | 6 | 10.3 | 21 | 18.1 | 0.5 (0.2-1.5) | 0.6 (0.2-1.7) | 0.366 |
| *Occupation involves forest visit | 20 | 34.5 | 12 | 10.3 | 4.6 (2.0- 10.2) | 3.8 (1.6-9.0) | 0.003 |
| History of travel | 9 | 18.4 | 15 | 12.9 | 1.2 (0.5-3.0) | 1.1 (0.4-2.9) | 0.916 |
| **B. Factors related to indoor dwelling environment** |  |  |  |  |  |  |  |
| *Other roofing materials | 10 | 8.6 | 3 | 2.7 | 7.9 (1.9- 45.7) | 13.6 (2.5-74.7) | 0.003 |
| Living in a house with cracked walls | 48 | 82.8 | 79 | 68.1 | 2.3 (1.0-4.9) | 3.9 (1.1-13.6) | 0.032 |
| Cracked floor type | 49 | 84.5 | 94 | 81.0 | 1.3 (0.5-3.0) | 1.4 (0.3-7.1) | 0.707 |
| 5 and below household occupants | 37 | 63.8 | 60 | 51.7 | 1.6 (0.9- 3.1) | 1.8 (0.8-3.8) | 0.131 |
| Sharing residence with household member with ulcerating disease | 16 | 27.6 | 3 | 2.3 | 14.4 (3.8- 79.3) | 16.0 (4.1-62.8) | 0.000 |
| **C. Factors related to outdoor environment** |  |  |  |  |  |  |  |
| Sighting rock hyrax near residence | 51 | 87.9 | 67 | 57.8 | 5.3 (2.2- 12.7) | 3.0 (0.9-9.3) | 0.065 |
| Sighting wild jackal near residence | 11 | 19.0 | 19 | 16.4 | 1.2 (0.5-2.7) | 1.2 (0.4-3.6) | 0.683 |
| Sighting porcupine near residence | 43 | 74.1 | 79 | 68.1 | 1.3 (0.7-2.8) | 0.5 (0.2-1.3) | 0.159 |
| Sighting mongoose near residence | 44 | 75.9 | 80 | 69.0 | 1.4 (0.7-2.9) | 1.1 (0.4-2.7) | 0.846 |
| Domestic dogs in the residence | 36 | 62.1 | 83 | 71.6 | 0.7 (0.3-1.3) | 1.3 (0.5-3.2) | 0.553 |
| Cultivated crop farm near residence | 46 | 79.3 | 113 | 97.4 | 0.1 (0.0- 0.4) | 0.1 (0.0-0.5) | 0.006 |
| Nearby forest or thicket | 53 | 91.4 | 67 | 57.8 | 7.8 (2.8- 26.4) | 7.0 (2.0-24.7) | 0.003 |
| Nearby open water source | 5 | 8.6 | 24 | 20.7 | 0.4 (0.1- 1.1) | 0.4 (0.1-1.1) | 0.081 |
| Immediate neighbour with ulcerating disease | 21 | 36.2 | 9 | 7.8 | 6.8 (2.8- 16.0) | 3.1 (1.1-8.8) | 0.031 |
| Distant neighbour with ulcerating disease | 17 | 29.3 | 41 | 35.3 | 0.8 (0.4-1.5) | 1.0 (0.4-2.4) | 0.937 |
| Presence of garbage mound near residence |  |  |  |  | 0.6 (0.3-1.1) | 0.6 (0.2-1.5) | 0.278 |
| *Other roofing materials- Grass, leaves, earthen, wood or rock (cave) |  |  |  |  |  |  |  |
| Cracked floors included earthen, rock (caves), or floors with visible crevices |  |  |  |  |  |  |  |
| Cracked wall type included earthen walls, rock (caves) or walls with visible cracks and crevices |  |  |  |  |  |  |  |
| Contact was defined as sharing a residence or living close to a neighbour within 150 meters or outside 150 meters radius as estimated by the enumerators |  |  |  |  |  |  |  |
| Water sources included river, dam, well, bore hole, pond or spring |  |  |  |  |  |  |  |

In the first category, individuals who preferred staying outside their residence in the evening after sunset (OR 4.1, CI 1.2-16.2) and those whose primary occupation involved visiting forests (OR 4.56, CI 2.04-10.22) had significant associations with disease in the bivariate analysis. After adjusting for other factors in the multivariate model, only occupations involving forest visit remained significant with a reduced odds ratio of 3.8. Activities such as charcoal burning, hunting, herding, stone masonry and mining were included among the occupations involving forest visits. When assessed separately, these occupations had significantly large odds ratios, but this analysis is not reported here due to possible close link between each of these occupations with forest visits. Other individual attributes such as sex, level of education, history of travel or use of mosquito nets did not have any significant association with disease. **Table 3.**

Three of the five factors that were fitted in the second category of factors related to indoor transmission remained statistically significant in the bivariate analysis. This included sharing same residence with a household member with typical skin lesions (OR 14.4, CI 3.8-79.3), residing in a house with alternative roofing materials (OR 7.9, CI 1.9-45.7) and residing in a house with cracked walls (OR 2.3, CI 1.0-4.9). These factors also showed significant associations with increased odds ratios after allowing for other factors in the multivariate model. **Table 3**

In the third category of factors related to outdoor environment, four of the eleven factors included in the analysis contributed significantly in this ‘group model’: sighting rock hyraxes near residence (OR 5.3, CI 2.2-12.7), residing near a forest (OR 7.8, CI 2.8-26.4) and living close to a neighbour with typical skin lesion (OR 6.8, CI 2.8-16.0) had increased likelihood of cutaneous leishmaniasis. These three factors remained significant in the multivariate model but with reduced odds ratios. Having a cultivated crop farm surrounding the residence (OR 0.1, CI 0.0-0.4) appeared protective. This association remained significant in the multivariate model. **Table 3**

In the final model, seven factors remained significant after controlling for all factors. Among factors related to the individual, occupations that involve visit to the forest remained significant in the final model (aOR 3.4, CI 1.1-10.7). Among the indoor factors, living in a house with cracked walls (aOR 5.5, CI 1.6-19.3), five or fewer household occupants (aOR 2.8, CI 1.1- 7.1) and sharing residence with a household member with typical ulcerating disease (aOR 26.7, CI 5.2-135.8) remained significant. Location of a forest in the neighbourhood of residence (aOR 5.8, CI 1.7-19.6) and having a neighbour with typical ulcerating disease (aOR 5.3, CI 1.8-15.7) remained significant among factors related to outdoor living environment. Having a cultivated crop farm near the residence (aOR 0.1, CI 0.0-0.5) remained protective. The odds ratios for these associations are as shown in **Table 4**.

**Table 4:** Final model showing adjusted odds ratios and 95% confidence intervals for factors related to cutaneous leishmaniasis in Gilgil, Kenya.

| Variable | Adjusted odds Ratio | 95% CI | P-value |
| --- | --- | --- | --- |
| *Occupation involves forest visit | 3.4 | 1.1-10.7 | 0.038 |
| Living in a house with cracked walls | 5.5 | 1.6-19.3 | 0.008 |
| Cultivated crop farm near residence | 0.1 | 0.0-0.5 | 0.007 |
| Nearby forest or thicket | 5.8 | 1.7-19.6 | 0.005 |
| Immediate neighbor with ulcerating disease | 5.3 | 1.8-15.7 | 0.003 |
| Sharing residence with household member with ulcerating disease | 26.7 | 5.2-135.8 | <0.001 |
| 5 and below household occupants | 2.8 | 1.1-7.1 | 0.029 |

## Discussion

This study has highlighted the burden of cutaneous leishmaniasis (CL) in Gilgil through records review and house-to-house survey. The burden of CL depicted in the records review could be higher. Our observation of quality of hospital records revealed gaps in standard surveillance case description and diagnosis of CL. The official health reporting portal of Kenya Ministry of Health through the DHIS2 platform does not incorporate cutaneous leishmaniasis among the monthly reports sent from health facilities across the country (20). In this portal, reports of CL will ordinarily be lumped and reported as “diseases of the skin” meaning that these cases never get to be reported at the national level. Coupled with inadequate diagnosis at health facility level, there is potential misdiagnosis and underreporting of CL.

Through analysis of potential demographic, behavioural and environmental attributes, we have been able to identify independent risk factors that are associated with cutaneous leishmaniasis in the study area. Significant associations seen in analysis of risk factors in this study suggest an overlap of factors that promote the likelihood of occurrence of CL at individual, indoor or outdoor levels. Individual factors included behavioural (spending time outside residence in the evening after sunset) and occupational factors (involving visit to the forest). Indoor risk factors included houses with cracked walls, households with fewer inhabitants and households with at least one infected member. Increased outdoor transmission was observed among individuals residing close to a forest, close to an infected neighbour or individuals who sighted wild rodents such as the rock hyrax near their residence.

The sand-fly vector for cutaneous leishmaniasis is described in literature as naturally anthropophilic and crepuscular, preferring to bite both indoors and outdoors in the evenings and early mornings (4,12,21). In previous studies, the sand-fly has been identified in parts on Kenya, including Gilgil (12,22–24). Among the indoor factors, we observed almost four-fold increased likelihood of CL among those who resided in houses with cracked walls. It is possible that large cracks and crevices on walls of residences could form perfect daytime hiding places for the sand-fly after a blood meal. Perhaps a more direct evidence of indoor transmission is supported by our finding of increased risk of CL in households where at least one member had typical ulcers (aOR 16.0, CI 4.1-62.8). This observation could be explained in part by familial clustering tendencies of CL that have been reported in a previous studies (25–27) and by a possible presence of high sand-fly vector density within these residential units (4).

Our finding of increased risk of CL in households with 5 or less inhabitants marks a departure from what has been observed in most studies since a large house-hold size (number of regular residents of a household) and high population density is considered a proxy indicators of poverty which has been associated with CL (2). One possible explanation for this relationship could be that small household sizes could predispose the few household occupants to potentially frequent sand-fly-human contact through bites (4). This relationship has been well studied in the case of other arthropod-vector borne diseases like malaria (28,29).

Presence of forest near residences and residents engaged in occupations that involved visit to the forests including charcoal burning, herding, hunting, wild honey harvesting and stone mining, all had increased likelihood of CL. Forest and thickets are likely to provide suitable habitats to immature and mature forms of the sand-fly vector. Additionally, the rocky caves and thickets around Gilgil have also been known to be infested with hyraxes and other mammals that are natural reservoirs of cutaneous leishmaniasis, and were likely to be sighted around homes among those with CL (9,30). As a result of increased population pressure, changes in land use (from forestry to agriculture) and deforestation, infected wild rodents have been known to migrate closer to human residences (2,4,21). Therefore, one plausible explanation for increased risk observed among residents who frequent forests for livelihood could be that such exposures would increase the frequency and duration of sand-fly-human contact. It appears that frequent vector-human contact either due to increased density of sand-fly in the human dwelling environment or as a result of humans venturing in sand-fly infested habitats in the forests could explain the increased risk of transmission in both indoor and outdoor settings, with the constant being increased exposure to the vector.

In our study, CL cases were clustered around two definite foci in Gilgil: most cases originated either from *Gitare*, the northern part of Gilgil along the rocky cliffs on the wall of the Great Rift Valley or from the southern end of Gilgil in *Útut* forest which is located in an area of solidified volcanic larva on the floor of the Great Rift Valley. Based on observations by the study team, both regions are remote, largely inaccessible, lack basic infrastructure and are inhabited by small scale farmers (*Gitare*) or forest dwellers (*Útut*). In *Gitare*, crop farmers were observed to be continuously encroaching the thickets around rocky escarpments near their homes for farmland and many new homes were seen to be constructed into areas that were previously under forest cover. In *Útut*, charcoal burning, stone mining, bee-keeping and herding are majorly carried out in the forests by a majority of residents.

The finding of clustering of cases in these two foci could indicate accelerated CL transmission in these areas. Indeed, Sang et al described *Utut* forest, among other regions, as a focus of CL on the floor of the rift valley with the sand-flies also identified in this region (9). This focalized transmission pattern could be attributed to three explanations. First, the sand-fly vector implicated in transmission of CL has a restricted flight radius, mostly flying within a range of 50 meters around their habitat (24). Secondly, there is abundance of mammal reservoirs in these localities as evidenced by the three-fold likelihood of sighting of rock hyraxes by the majority of CL patients in the study area. Wild rodents including rock hyraxes have been described in literature as possible animal reservoirs of CL (9). Lastly and importantly, these two localities epitomize high rate of human encroachment to previously non-inhabited land. In *Gitare*, crop farming is expanding while in *Útut* forest, more people are visiting the forest to burn charcoal, harvest honey, mine stones and herd livestock.

Studies have consistently shown more cases of CL among poor, neglected populations who are likely to be less educated and mostly unemployed (2,31). In our study, the majority of the sampled population (90.8%) had less than primary level of education, 20% were unemployed and relied on menial jobs to occupy their time and generate income and a 72.1% used herbal medication for treatment of skin lesions. All these findings reinforce the above assertion. Other publications have also shown similar findings (32,18). However, analysis of level of education and employment status did not show significant associations in the bivariate analysis in our study.

As expected, cases were either younger or older individuals. This finding is consistent with available literature that has shown that young age and immune-suppression associated with old age are significant risk factors to cutaneous leishmaniasis (32–34). A possible explanation for this observation could be due to immune dynamics of the agent; the younger cases are immune naïve hence more vulnerable to primary infection while the older persons could be susceptible to reactivation of old infections or increased risk of primary infection. However, the explanation for increased infections among younger persons may not be supported by another finding in our study that some cases had both active lesion (typical ulcer) and old scars at various stages of healing. This could imply repeated infections in the same individual, a possible indication that immunity developed following primary infection could be short-lived as seen elsewhere in literature (35).

The most common presentation seen among the cases in this study was skin ulcers frequently located in head and neck regions of the body. Other lesion locations were hands and feet. Similar findings have been recorded in other studies (9,33). The wounds mainly affected exposed areas of the body which are commonly bitten by the sand-fly. The mouth and nostril ulcers observed among the cases, could point at a mixed form of leishmaniasis with a possibility both the cutaneous and muco-cutaneous forms of the disease in the area. Further clinical evaluations would be necessary among the cases to determine if both forms of this disease affect these residents.

Despite its strengths, this study had some limitations. Most hospital records available for the review were either incomplete or inaccurate. Lack of a proper hospital records could lead to underestimating the burden of cutaneous leishmaniasis. It was also not possible to compute other parameters such as incidence of disease as some of the cases enlisted reported multiple infections over time. Moreover, cutaneous leishmaniasis has a long latency period hence some cases identified in the hospital records could be prevalent cases as opposed to incident cases. Despite our finding of strong association between individual, indoor and peri-domestic factors in the transmission of CL, our study did not estimate the relative contribution of each of these factors in transmission of CL.

### Conclusion and recommendations

The burden of cutaneous leishmaniasis in Gilgil is large. However, due to sparse data in the visited facilities, the true burden of disease could be higher. Cases were reported throughout the years, consistent with a locally endemic disease with periods of increased cases reported suggesting possible seasonal pattern of transmission. There is strong evidence for both indoor and outdoor patterns of disease transmission as suggested by clustering of cases among household members and focalized transmission pattern in specific neighbourhoods. Occupations and activities that involve visiting forests or residing near forests and residing in a house or neighbourhood with a person with CL were identified as significant exposures of the disease. CL lesions mostly affected younger or older exposed residents in exposed parts of the body.

There is need to institute and strengthen surveillance for cutaneous leishmaniasis in Gilgil in order to provide a better estimate of the disease burden. Emphasis should be put on data quality in the affected health facilities. Focalized transmission of disease driven by human interaction with known sand-fly habitats such as forests points at possibility of success of tailored control interventions like indoor residual spraying and insecticide treated nets. We recommend that further studies be conducted to establish effectiveness of adequate sand-fly bite protection e.g. by wearing protective clothing among the residents of Gilgil. Modification of human behaviour in areas of known transmission risks would appear to be potentially effective and feasible as a disease control strategy. Discouraging practices such as hunting and charcoal burning would be in theory be effective but would not be practical. However, the role of environmental factors and wild mammals in disease transmission should be investigated further.

## Acknowledgements

I would like to acknowledge the contribution of the following individuals and institutions for their various contributions in this work: Dr Joe Lenai and the Nakuru County Health Management Team, Dr. Ester Kanyiru and the entire Gigil Sub County Health Management team, International Livestock Research Institute (Kenya), FELTP Faculty (Kenya), Washington State University Global Health programs Kenya (WSU-GH Kenya) and the Unit for Neglected Tropical Diseases in Kenya. The findings and conclusions in this paper are those of the author and do not necessarily reflect the official views of the supporting institutions.

## Supporting information legends

**Figure 1:** Map of the Investigation area, Gilgil, Kenya 2016

**Figure 2:** Flow diagram of selection of cases and controls before and after field work

**Figure 3:** Trends of patients treated for suspected cutaneous leishmaniasis in health facilities

**S1 Checklist:** STROBE Checklist

